# Quenching an active swarm: Effects of light exposure on collective motility in swarming *Serratia marcescens* colonies

**DOI:** 10.1101/331801

**Authors:** Alison E Patteson, Junyi Yang, Paulo E Arratia, Arvind Gopinath

## Abstract

Swarming colonies of the light responsive bacteria *Serratia marcescens* grown on agar exhibit robust fluctuating large-scale collective flows that include arrayed vortices, jets, and sinuous streamers. We study the immobilization and quenching of these large-scale flows when the moving swarm is exposed to light with a substantial ultra-violet component. We map the response to light in terms of two independent parameters - the light intensity and duration of exposure and identify the conditions under which mobility is affected significantly. For small exposure times and/or low intensities, we find collective mobility to be negligibly affected. Increasing exposure times and/or intensity to higher values temporarily suppresses collective mobility. Terminating exposure allows bacteria regain motility and eventually reestablish large scale flows. For long exposure times or at high intensities, exposed bacteria become paralyzed, with macroscopic speeds eventually reducing to zero. In this process, they form highly aligned, jammed domains. Individual domains eventually coalesce into a large macroscopic domain with mean radial extent growing as the square root of exposure time. Post exposure, active bacteria dislodge exposed bacteria from these jammed configurations; initial dissolution rates are found to be strongly dependent on duration of exposure suggesting that caging effects are substantial at higher exposure times. Based on our experimental observations, we propose a minimal Brownian dynamics model to examine the escape of exposed bacteria from the region of exposure. Our results complement studies on planktonic bacteria and inform models for pattern formation in gradated illumination.

## 1 Introduction

Swarming motility is a flagella driven mode of bacterial surface migration that is widespread in both gram-positive and gram-negative bacteria and enables rapid colonization of environments ^1–8^. Swarming is typically initiated when free-swimming (planktonic) bacteria grown in fluids are then transferred to soft wet agar gels^5^. The transfer triggers a change in phenotype; individual cells become significantly elongated and the number of flagella increases, sometimes to 10–100^17^. At high densities, the highly mobile expanding colony develops complex, long-range intermittent collective flow features that involving multiple bacteria traveling in rafts or flock-like clusters. Speeds are usually high near the edge of the expanding colony with intensity (magnitudes) decreasing away from the propagating edge^2,4,5,9^ and towards of center of the colony. Recent studies^10–12^ demonstrate that swarming confers multiple benefits including enhanced colonization rates and elevated resistance to antibiotics. Swarming is also found to be co-regulated with virulence, and also implicated in infectiousness and fitness of pathogenic bacterial species^3,7^.

Healthy bacterial cells - in both planktonic as well as collectively moving states - sense spatiotemporally distributed cues, continuously process these inputs and transduce them to variations in motility and other responses^1,4,9,13^. For instance, single bacteria are observed to respond to chemical and mechanical stimuli by modulating and controlling the molecular motors underlying flagellar motion^14–18^. Intense light with wavelengths in the range 290-530 nm encompassing the ultraviolet (UV) range is known to trigger changes in the motility of planktonic chemotactic bacteria^19,19–21^. Prolonged exposure to high-intensity light results in progressively slow swimming with paralysis occurring eventually^19,20^ due to irreversible motor damage. Striking variations in motility associated with modulation in the functioning of the MotA - MotB pair and FliG comprising the rotary motor complex in the flagella - are also observed in chemotactic bacteria. The net result is a change in the swimming gait - specifically, tumble length and tumble frequency^15,17^ as for instance seen in *Escherichia coli* and *Streptococcus* ^20,22^. Connecting and relating these single cell responses to variations and changes in swarming bacterial systems however poses significant challenges and remains a topic of active research.

From a medical perspective, light treatments employing UV-A, UV-B and UV-C radiation are emerging as attractive alternatives to antibiotic treatment of pathogenic bacteria. Light exposure is known to inhibit cell growth and induce gene damage^23^ in marine organisms (alphaprotobacteria and bacterioplankton,^24^), airborne bacteria^25^, as well in bacterial biofilms^26,27^. Irradiation of surfaces using blue light and phototherapy that activates endogenous or exogenous photosensitizers^13,28,29^ has been shown to sterilize and disinfect bacteria laden surfaces and swarming bacteria. Intense light is also known to promote wound healing^29,31^ with visible light recently approved to treat bacterial infections such as acne^32^. Given these promising studies and the timeliness of light treatments, understanding the connection between motility, infectiousness and light exposure is particularly important.

Here, we report on the effects of wide-spectrum light with significant UV components on the collective motility of swarming *Serratia marcesens*. In collectively-moving swarms, individual self-propelling cells are influenced by steric and hydrodynamic interactions with their neighbors^4–6,33–35^. These interactions result in complex features including fluctuating regions of high vorticity and streamers. We first discuss statistics of our base state - that is features of collective motility in unexposed bacterial swarms. Following this, we report and analyze the drastic variations in mobility evident when localized regions of the swarm are exposed to wide-spectrum light. As a part of this process, using two orthogonal parameters - the light intensity and duration of exposure - we identify the conditions under which mobility is affected significantly. Interestingly, this *quenching* of activity may happen in either reversible or irreversible manner. For low intensities or short exposure times, bacteria recover their motility, reestablish collective flows, and erode the passive domain from within when exposure is terminated. For long exposures or high intensity light, actively generated collective motion ceases slowly and the passive region grows with time until saturation due to the finite extent of the light field. As this quenching progresses, dense strongly jammed domains of immobile cells form and hinder the penetration of unexposed bacteria into the region, thereby localizing the damage in the colony. Post-exposure, swarming cells penetrate into the previously passive domain; associated emergent flows dislodge and then convect immobile bacteria away with the initial dissolution rate dependent on the duration of light exposure. We hypothesize that this dependence comes from the highly jammed and aligned configurations of the paralyzed bacteria attained for long exposure times.

To complement our experiments and to investigate how subpopulations of exposed bacteria may be able to extricate themselves and escape the light, we propose and analyze a minimal agent based Brownian dynamics model for a illuminated test cell. By incorporating the effects of light exposure through changes in the rotational and translational diffusivity of this model cell, we demonstrate that variations in bacterial spreading distances may arise in a population comprising cells featuring a distribution of self-propulsion speeds. Subpopulations of exposed bacteria corresponding to the fastest moving cells may escape from the exposed region; significantly however, the dense packing in collective swarms may alleviate this effect suggesting that exposure time is as important as light intensity.

## 2 Experimental methods

Swarms of *Serratia marcesens* (strain ATCC 274, Manassas, VA) were grown on agar substrates, prepared by dissolving 1 wt% Bacto Tryptone, 0.5 wt% yeast extract, 0.5 wt% NaCl, and 0.6 wt% Bacto Agar in deionized water. Melted agar was poured into petri dishes, adding 2 wt% of glucose solution (25 wt%). The bacteria were then inoculated on solidified agar plates and incubated at 34°C. Colonies formed at the inoculation sites and grew outward on the agar substrate from the inoculation site.

These spreading swarms were studied and imaged 12-16 hours after inoculation. The bacteria were imaged with the free surface facing down with an inverted Nikon microscope Eclipse Ti-U using either a Nikon 10x (NA = 0.3) or 20x (NA = 0.45) objective. Images were gathered at either 30 frames per seconds (fps) with a Sony XCD-SX90 camera or at 60 fps with a Photron Fastcam SA1.1 camera. We used videos of the swarm and PIVLab software ^36^ to extract the velocity fields of the bacteria with particle image velocimetry (PIV) techniques. Particle image velocimetry determines velocity fields by calculating local spatial-correlations between successive images. Here, the images are of bacteria (either active or passive) such that PIV yields the bacterial velocity fields directly and not the velocity field of the ambient fluid. We sampled the velocity field at 3 *μ*m spatial intervals in the images, checking the frame rate for accurate resolution. All the analysis of the data extracted from the videos was done using freely available object-oriented Python software.

To mimic the exposure of bacteria to naturally occurring high intensity light, we used a wide spectrum mercury vapor lamp^37^. Standard microscope optical components were used to focus the light on the swarm. The bare unfiltered maximum intensity (measured at 535 nm) of the lamp *I*_0_ was reduced to lower, filtered (maximum) intensities *I* using neutral density filters. Both *I*_0_ and *I* depend on the objective and this effect was taken into account in all experiments by calibrating for the exposure intensity. Using a spectrophotometer (Thorlabs, PM100D), we measured maximum intensities *I*_0_ = 980 mW/cm^2^ (at 535 nm) for the 10x and *I*_0_ *=* 3100 mW/cm^2^ (at 535 nm) for the 20x objectives. Note that the actual intensity of light *I*\* (**r**, *t)* as measured in the swarm is spatiotemporally varying due to bacterial motion and also due to the associated point spread distribution as the light passes through the aperture. Typically, the maximum intensity corresponds to that measured at the center (**r** = **0**) of the focused light beam.

## 3 Results and Discussion

Figure 1 is a snapshot of the expanding colony, a region of which (white area) is exposed to high-intensity light from a mercury vapor lamp^37^ using an octagonal aperture. The swarm is expanding from left to right; the colony edge is indicated in white in the figure. Bacteria swim in a thin layer above the agar substrate (inset, Figure 1, see also SI Movie 1). While we are unable to directly quantify the thickness of the dwarming layer, we expect the thickness of the swarm to vary with distance from the leading edge. Studies on *E*. *coli*^38^ swarming on agar substrates showed that cells form a monolayer over a significant region close to the leading edge, beyond this cells can form multi-layered regions. The same features can be anticipated on the experiments described here as well; consequently, we choose to illuminate the region of the swarm that is within the thin region.

**Fig. 1.**
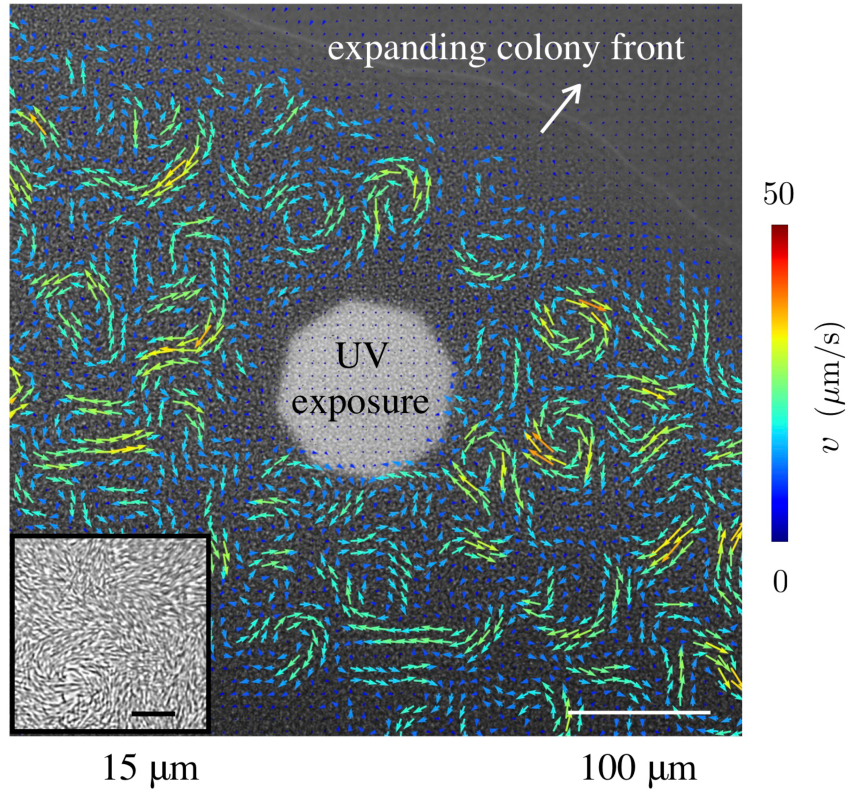
Characteristics of swarming and expanding colony: Snapshot of a *Serratia marcescens* colony on an agar substrate, during exposure to high-intensity light from a mercury lamp source; PIV derived velocity fields are overlaid in color. Swarming motion is pronounced approximately 50 microns from the expanding colony front. The inset shows pre-exposure bacterial alignment and density 150 *μ*m from the colony front.

Overlaid on the image in Fig. 1 are bacterial velocity fields gathered from PIV; the image is taken after 80 seconds of exposure. Outside of the exposed region, the velocity field exhibits long-range collective flows. Unexposed bacteria move fastest approximately 100-400 *μ*m from the colony edge (maximum speeds ∼80 *μ*m/s, average speed ∼30 *μ*m/s). In contrast, inside the exposed region, the bacterial motility is significantly impaired. This is evident in SI Movie 2; as swarming bacteria are exposed to light, they slow down and are eventually trapped within the exposed region. This feature is reflected in the trajectories of the small tracer-like particles as they slow down and eventually stop moving when trapped amongst the passive bacteria. The interphase boundary between the unexposed (passive) domain and the unexposed (active) part of the swarm features strong vortices, jets, and streamers, extending up to just a few microns away from the exposed domain.

### 3.1 Collective motility: Phase-space for reversible and irreversible quenching

The quenching (passivation) of collective mobility of the initially active bacteria is not immediate and can be reversible or irreversible depending on the intensity of light and the duration of exposure suggesting that the net dosage of light controls the response. This is consistent with previous studies at the single bacteria level. Based on these observations, we explore two features of the response to high intensity light: (i) the reversible versus irreversible nature of the bacterial response and (ii) the effects of exposure time and light intensities on the passive domain growth rate during exposure and dissolution rate post-exposure.

Figure 2 illustrates the three modes of bacterial response to light; our classification is based on averages obtained from time-dependent swarm velocity fields: exposed cells either (i) retain mobility with negligible effects, (ii) transiently stop moving and then regain motility at the single cell and collective level (reversible), or (iii) permanently stop moving (irreversible). To quantify the response, the behavior was mapped onto a phase diagram with exposure time *τ* and light intensity *I* (varied from the bare value *I*_0_ using neutral density filters) as variables. For sufficiently small exposure times (*τ* ∼ 20 — 40 s) and intensities (*I* < 220 mW, at 535 nm), exposed cells remain always active. Conversely, for large exposure times (*τ* > 60 s) and sufficiently high intensities (*I* > 220mW, at 535 nm), the bacteria are permanently passive (over the duration of the experiment). As seen in Fig. 2(a), between these two phases lies the temporarily passive, reversible case.

**Fig. 2.**
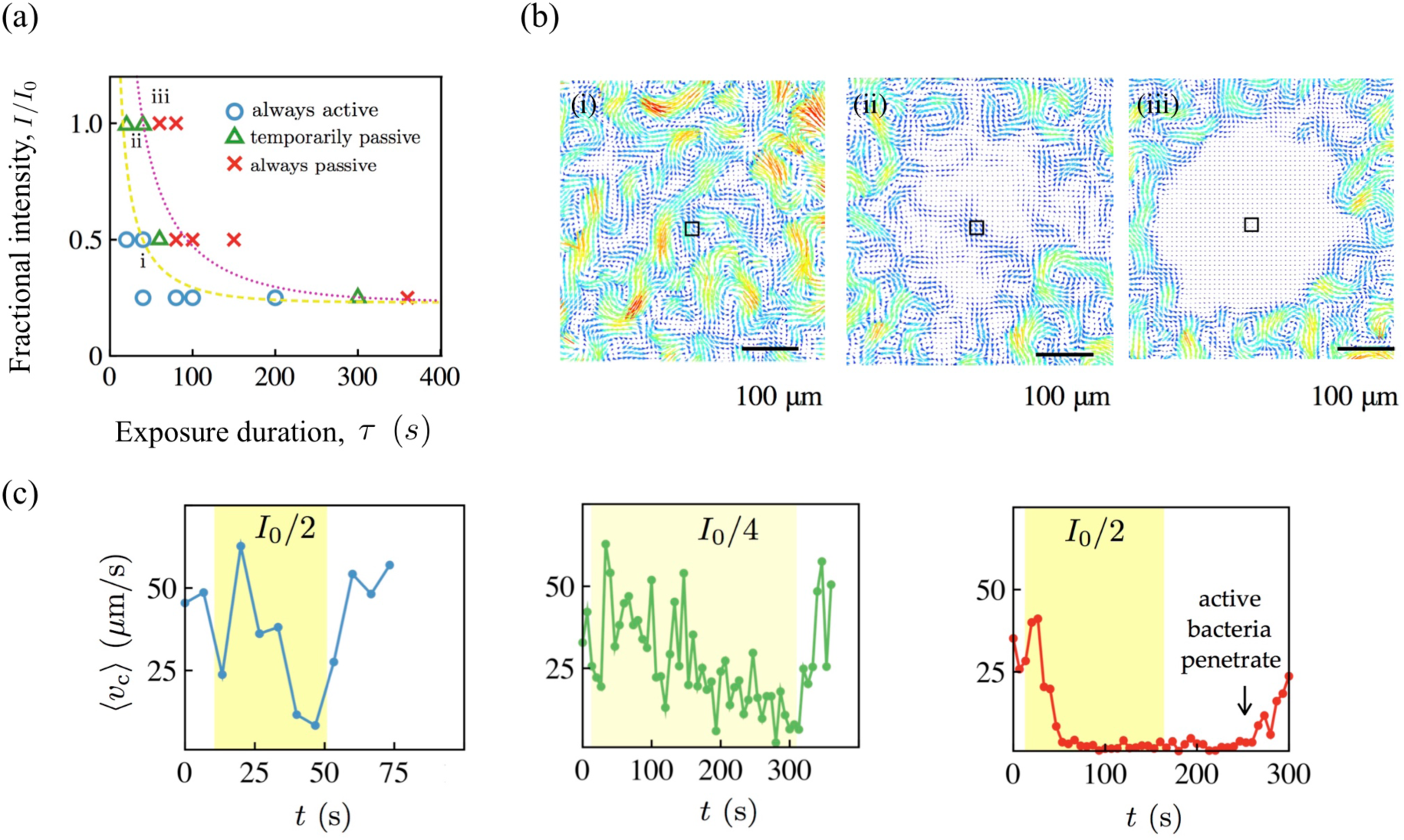
Phase space for collective motility after exposure: (a) Changes in collective flows (relative to the unexposed state) in swarming *Serratia marcescens* depend strongly on intensity and duration of light exposure. We use a wide spectrum mercury lamp (bare intensity *I*_0_ = 980 mW/cm^2^ at 535 nm) with filters to selectively expose regions of the swarm to a filtered maximum intensity I. From subsequent PIV analysis of bacterial velocities, the response can be classified into one of three types - (I) always active, (II) temporarily passive, and (III) always passive. The yellow (dashed) and pink (dotted) curves are phase boundaries predicted by equation 1. (b) Velocity fields taken 10 s post exposure are shown for each phase. Colors reflect speed with the arrows denoting polar orientation. Collective motility of temporarily immobile bacteria is recovered in approximately 15 seconds past exposure. (c) We plot the average speed of the swarm in the central region, highlighted by the box in (b) for times encompassing pre-exposure, exposure (yellow band), and post-exposure. Pre-exposure, the average swarm speed fluctuates between 25 to 50 *μ*m/s. For case (I), the bacteria briefly slow down during exposure, but recover in 6 s. In (II), the swarm speed approaches zero during exposure and recovers in 12s. In (III), the collective swarm speed drops to and remains zero.

The differences between the three (collective) motility regimes are highlighted in Fig. 2(b) and (c); these show PIV-derived velocity fields taken 10 seconds past exposure (Fig 2b) and the average bacterial speed ⟨*v*_c_⟩ - in the exposed region - over time (Fig 2c). Here, ⟨*v*_c_⟩ is the average speed of the velocity fields in a 22×22 *μ*m^2^ area, located at the center of the exposed region. In case (i), exposed cells remain motile and continue to exhibit long-range collective motions; the speed decreases but does not fully reach zero during exposure. The speed recovers to pre-exposure levels approximately 5 seconds after exposure. In case (ii), bacteria stop moving during exposure, yet spontaneously start moving again ∼ 1-10 s after the light is switched off. The cell speed ⟨*v*_c_⟩ takes longer to recovers to pre-exposure levels than case (i), and the recovery occurs heterogeneously with cells moving within the temporally-quiescent region (Fig. 2(b)-ii). This process superficially resembles the heterogeneous, disconnected melting of a large frozen domain. In case (iii), cells stop moving during exposure and do not regain their motility afterward for the whole duration of the experiment (20-300 s). Unlike case (ii), cells in the exposed region here do not spontaneously regain their motility (Fig. 2(b)-iii). We find that, in this case, the passive domain evolves solely due to its interaction with the active swarm at its boundary. The active swarm convects passive bacteria away from the boundary and the passive phase is dismantled entirely; the speed ⟨*v*_c_⟩ eventually increases (Fig. 2(c)-iii) as the swarm recaptures the quenched area.

Similar responses have been observed earlier in studies on *E*. *coli* and *S*. *typhimurium*^19^. Specifically, prolonged exposure to unfiltered light in both bacterial species resulted in constant tumbling, and eventual paralysis. *S*. *typhimurium* was found to responds instantaneously to exposure with recovery of normal motility in 2 s or less upon cessation of exposure provided the duration of exposure was less than 5s. Similarly, *E*. *coli* recovered normal motility in 1 to 10 s after cessation of exposure. Sustained exposure that ultimately results in paralysis was however found to be irreversible (with no recovery of motility even after 15 minutes) for both species.

The regimes in the phase-plot Figure 2(a) can be rationalized by assuming the existence of a lower threshold for the intensity *I*_min_ below which bacteria are not affected. We note that the curves separating these regions of phase space do not correspond to a single or unique value of the light dosage (intensity multiplied by exposure time) or net power. The response to light involves changes to the motor complex and possibly involves a cascade of biochemical events. At the same time, while active swarming bacteria swim in and out of the exposed region close to the edge; in the interior of the exposed region they are caged in by their neighbors. To interpret Figure 2, we treat these independent motility affecting mechanisms in an approximate phenomenological way using a lumped time approximation. We invoke an intrinsic time scale *τ*\* that determines the internal organismal response to light (here quantified using the maximum of the filtered intensity) resulting in either temporary (*τ*\* = *τ*_temp_) or permanent (*τ*\* = *τ*_perm_) passivation. The curves that separate the different responses in Figure 2(a) may then be fit by

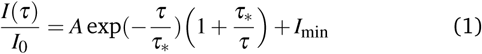

where the constants (*A,I*_min_, *τ*_temp_) are (0.2,0.23,62) for the yellow (dashed) curve and (*A,I*_min_, *τ*_perm_) are (0.33,0.23,100) for the pink (dotted) curve. The bare (unfiltered, maximum) intensity *I*_0_ = 980 mW/cm^2^ at 535 nm. As expected, *τ*_temp_ < *τ*_perm_.

The recovery of collective motility when sustained exposure is not maintained has significant implications. In patterned or localized light fields, fast cells have a higher chance of escaping the exposure region prior to complete paralysis. While swarming bacteria can reorient by run and tumble movements, motility driven by close bacteria-bacteria interactions dominate in a swarm. Thus slower cells are impacted more; first, because of longer exposure to the light and second, because they are more easily caged in and trapped by already paralyzed cells. In periodic fields, these persistence bacteria may also completely recover by the time they encounter the next exposed region.

### 3.2 Form and growth of the quenched region

To determine the extent of the quenched passivated domain, we use two threshold-based methods: the first is based on analysis of image intensity fluctuations^40,41^ and the second utilizes PIV derived velocity fields. Our experiments were for the most part (unless stated otherwise) conducted at a filtered intensity of 3100 mW/cm^2^ (measured at 535 nm^37^); exposure times *τ* were varied from 10-300 seconds.

The first method to extract the boundary of the quenched region uses point-wise fluctuations of the spatially varying intensity

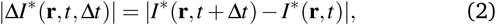

where *I*\* (**r**,*t*) is the two-dimensional image intensity at pixel position r and Δt is the time step. To reduce noise in the system (due to pixel resolution, short-range fluctuations, and background light fluctuations), we filter pixel-wise |Δ*I*\*(r,*t*, Δ*t*)| by smoothing over 3×3 *μ*m^2^ areas. We then varied Δt to obtain results that resulted in well resolved domains. We found that using Δt = 0.1 sec, which corresponds to the time in which a swarming *Serratia* cell swimming at 50 *μ*m/s moves roughly a body distance (5 *μ*m) provided good results; variations in At around this value (0.05 s < Δ*t* < 0.3 s) resulted in only small variations in results. As shown in Fig. 3, the intensity fluctuations allow us to clearly identify and distinguish two domains, the immobile domain where values of |Δ*I*\*| are relatively small and the motile domain where |Δ*I*\*| are relatively large. Thresholding |Δ*I*\* | then yields the locus of points that defines the boundary of the active and passive phases.

**Fig. 3.**
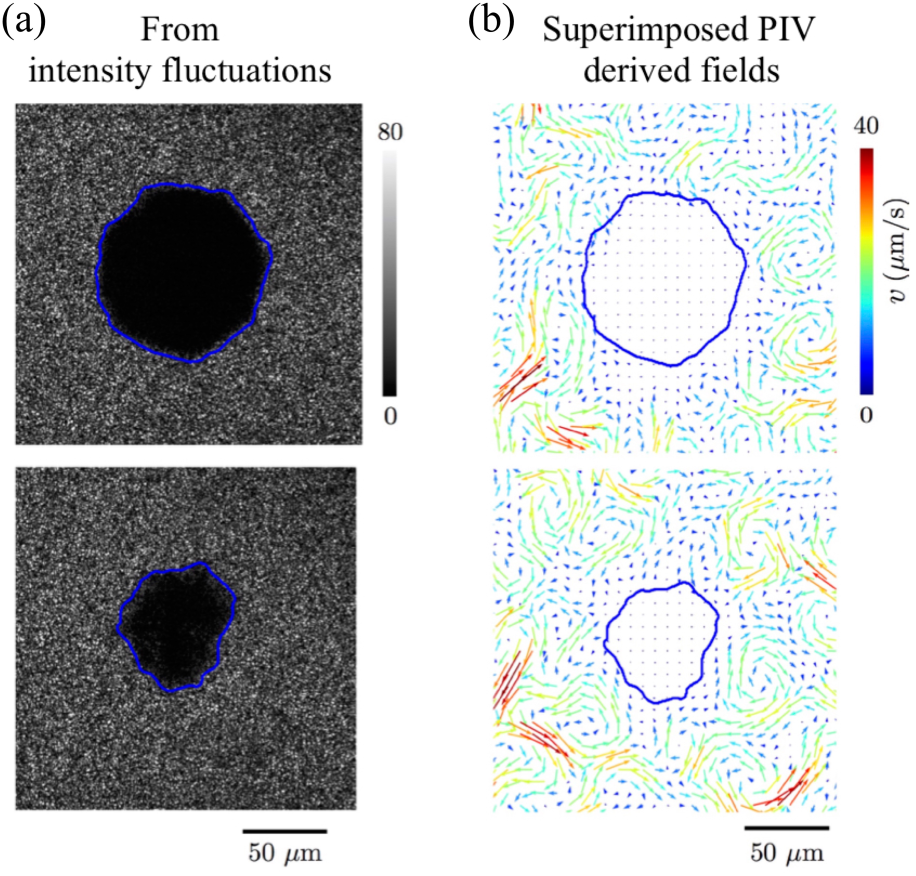
Thresholding using intensity fluctuation fields: (a) We calculate an intensity fluctuation map |*I*\*(**r**,*t* + Δ*t*) -*I*\*(**r**,*t*)|, with Δ*t* = 0.1 seconds. Intensity fluctuations are low (black, third tile from left) in the regions where the swarm is not moving. The map when thresholded allows us to identify the boundary position (blue contours). The passive phase shrinks as time goes by as shown at *τ* = 1 sec (top) and *t* = 40 sec (bottom) post-exposure (Exposure duration *τ =* 80 sec, Intensity is 3 W/cm^2^ measured at wavelength 535 nm). (b) PIV derived bacteria velocity fields confirm the boundary location obtained from the intensity fluctuation maps. A mathematically defined diffuse boundary may be obtained from phase field profiles^39^ using order parameters^33^; here however, we use simple thresholding.

In the second method, the boundary position was obtained from coarse-grained spatially-averaged bacterial velocity fields from PIV (Fig. 3(b)); again, the exact boundary was defined using a threshold criterion^33^. The locations of the active and passive domains, as well as their relative sizes such as area, match between the two methods, although the intensity fluctuations identifies smaller features of the boundary compared to that from PIV. These metrics capture complimentary aspects of the swarm’s motility: the intensity fluctuations are a scalar measure of density fluctuations and PIV yields vectorial velocity fields that quantify instantaneous polarity fields. In summary, simple thresholding using the intensity fluctuations and/or PIV derived velocity fields, yields physically meaningful boundary positions that separates the quenched region from the swarm. This approach can be formalized using phase-field approaches in order to extract the location and width of the interface^33,39^.

We next analyze the role of the exposure time *τ* that together with *I* determines the total light dosage, on the shape, size, and dissolution rate of the passive domain that is surrounded by the active swarm. Phenomenologically, exposure to light here is akin to quenching with activity *modified and in a sense removed* from our system. That is, exposure reduces the activity or energy in the system by removing the ability of bacteria to self-propel and inhibiting and eventually preventing large scale collective motions. Furthermore, as the bacteria slow down cell-cell steric interactions result in tightly jammed clusters with distinct aligned domains (Figure 3(a)).

In order to quantify additional features, we reduce variables to coarse-grained one-dimensional profiles. For instance, a typical measure of the bacterial speed ⟨v⟩ is obtained by averaging over the azimuthal angle. This yields a field that is a solely a function of the radial distance *r* from the center of the exposed region where we expect the light intensity to be a maximum.

When the light source is turned on, the bacterial speeds in the exposed region decreases non-uniformly, diminishing the most in the center of the exposed region (Fig. 4a). The two dimensional velocity fields (snapshots in time) suggest that bacteria stop moving in multiple sub-domains, each of which continuously interact with active bacteria that are still moving. It takes approximately 50 seconds of exposure for a spatially-uniform passive domain to develop. In the swarming state, the motion of individual *Serratia marcescens* may be decoupled into a mean velocity (with long range correlations arising from collective motions) and a diffusion-like term dependent on bacterial diffusivity and on steric cell-cell interactions. We hypothesize that the delay we observe time in going from the temporarily passive to fully immobile phase is a consequence of slow and differing time scales over which collective speeds and diffusivities decrease. Both these trends result in the slowly deactivating bacteria experiencing increasing jammed situations with local flocks aligning. Performing a radial average allows us to obtain a reduced map of the bacterial speeds ⟨v⟩ over time *τ* as shown in Fig 4(b) and 4(c). The exposed (quenched) domains appear as the dark region where the bacterial speeds are negligible. Once this initial quenched domain forms (after the initial lag time *t*_lag_), the size of the passive domain increases monotonically with time *t*. Fluctuations in the calculated extent of the quenched domain may arise physically due to variations in the (averaged) heterogeneous, fluctuating velocity fields during exposure (Fig. 4(a), tiles 2-5) that result in momentum boundary layers at the edge.

**Fig. 4.**
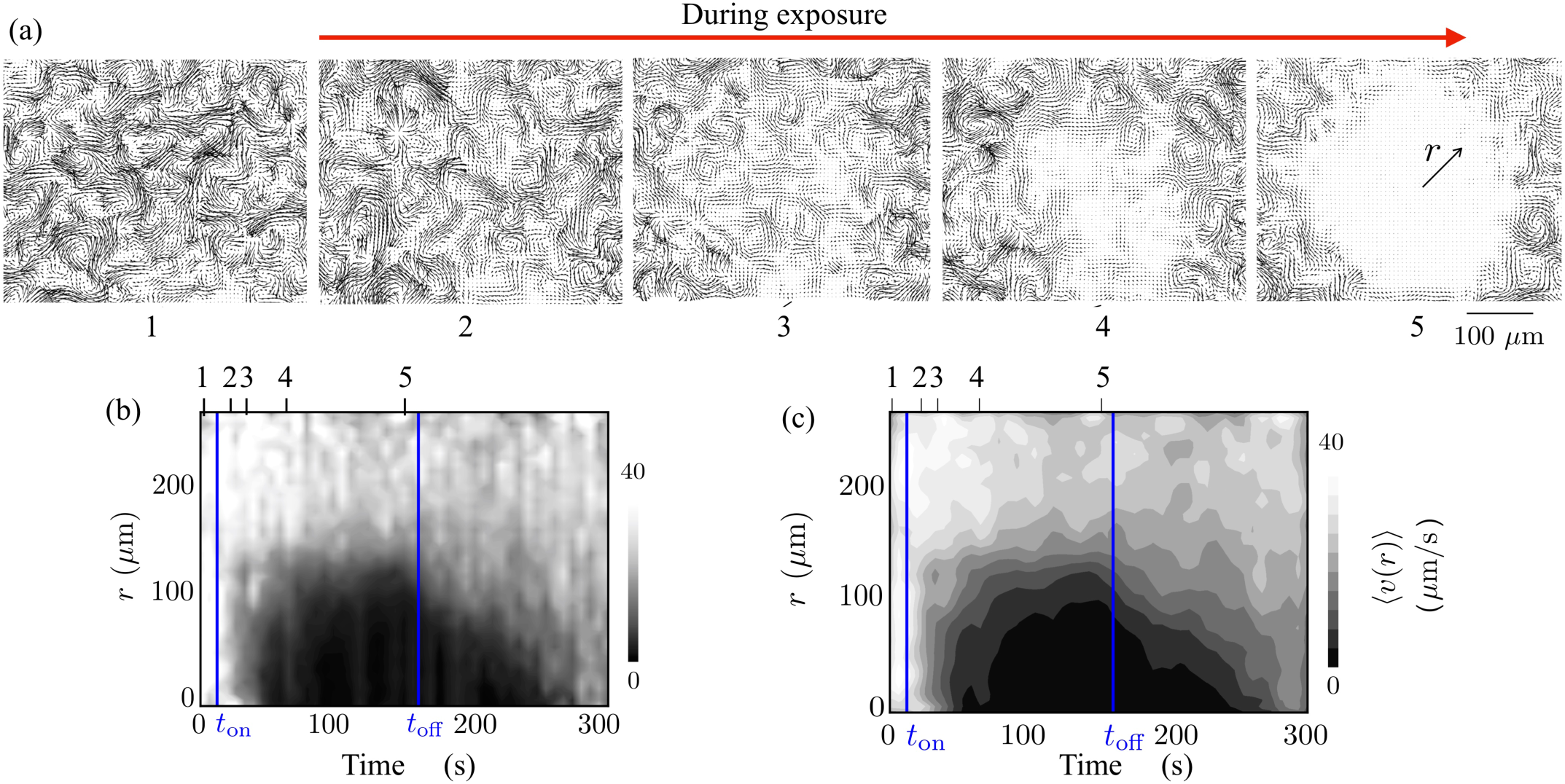
Thresholding using PIV derived fields allows for identification of edge of quenched region: (a) (Top) PIV derived bacterial velocity fields before (1) and during (2-5) light exposure. We note from plate (5) that as the exposure is continued, the paralyzing effects persist only a short distance into the active unexposed region as evident from the vortical structures seen near the edge. (b) The azimuthally averaged velocity ⟨*v*⟩ highlights the creation of an immobile quenched domain within the exposed region. When the light is switched off, the active bacteria from the unexposed regions penetrate into the quenched domain, eroding it away. The filtered maximum intensity = 500 mW/cm^2^. We note the brief lag after the light is switched on (at *t*_on_), the gradual increase to a finite size as *τ* ⟶ *t*_off_ and the rapid erosion and mixing with the grey interphase region *τ > t*_off_. The radial extent obtained by thresholding the intensity fluctuations also follows the square root dependence. (c) Increasing the area used to smoothen the intensity plots yields smoother but less sharp contours. We varied the smoothing box and Δ*t* so as to obtain convergent results.

When the light is turned off, active bacteria penetrate the passive phase and convect passive bacteria away. The advancing swarm intermingles with the passivated bacteria as it propagates inward. In approximately 100 seconds post exposure, the active swarm recaptures the exposed domain. We note that the boundary layer (grey region intermediate between the black and white regions) at the edge of the rapidly eroding quenched domain is larger than when the domain is being formed.

Note an (instantaneous) two dimensional boundary separating the exposed and unexposed regions may be obtained by using a cut-off speed for the two dimensional velocity field. From Fig 5(a) (aperture size 60 *μ*m), it is clear that the passivated domains that form can be irregular and asymmetric domains. As the exposure time is increased, the domains become larger (eventually comparable to the aperture size) and more regular. The maximum intensity of light is at the center of the aperture and so the quenching starts there with the boundary propagating outward. Nonetheless, as time progresses, the irregularity in the shape reduces as formerly separated mini-domains combine and form a single larger quenched domain. After this initial phase, the use of a one-dimensional speed field is justified. To obtain a quantitative estimate of the effective size of the quenched domain in Fig. 4(b); we threshold ⟨*v*⟩. Varying the thresholding value, we find an optimum of 10 *μ*m/s that yields physically meaningful results.

**Fig. 5.**
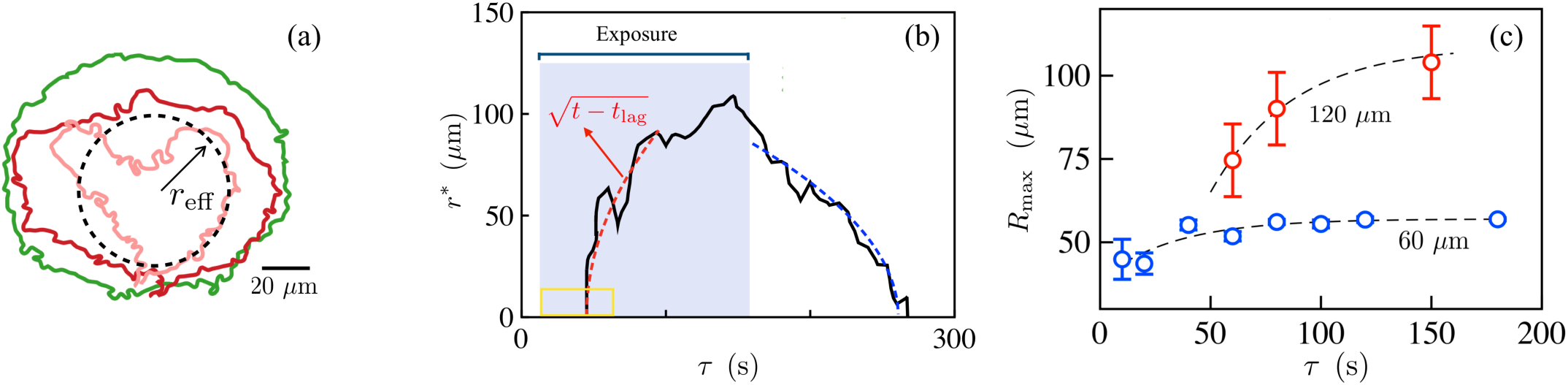
Growth, shape and dissolution of the quenched domain: (a) Interface shapes of the quenched (passive) region obtained from thresholding intensity fluctuation fields; we define *r*_eff_. Pink, red and green curves correspond to exposure times of 10, 20 and 100 seconds (aperture size 60 *μ*m, 20x objective). (b) The effective size of the quenched region grows during exposure, stabilizes while illuminated asymptotes to a constant and then decreases to zero once the light is switched off. We calculate the effective radius of the quenched region *r\** defined by the locus of points satisfying ⟨*v*⟩ (*r*\*) = 10,um/s and examine its dependence as a function of time. When the light source is turned on at *t*_on_, this radius increases from zero only after an apparent lag time *t*_lag_ *μ* 50s. Lowering the threshold velocity yields a noisier initial growth region with shorter lag. The initial growth has a square root dependence with time (red dashed curve). We observe deviations of around 5 – 10 *μ*m in these curves due to variations in the velocity field. The aperture size used here is 120 *μ*m. (c) The maximum effective size of the passive phase *R*_max_ increases with exposure duration *τ* and asymptotes to a constant that is less than the aperture size.

Note that this value is much less than the average speed of the active region (40 *μ*m/s) while large enough to average over small fluctuations.

Adjusting for the lag time, we find that the radius of the quenched region 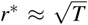 - shown in Fig 5(b) - where *T* = *t* − (*t*_lag_ + *t*_on_) with *t*_on_ being the time when the light source is turned on. Large variations in *r\** are a consequence of asymmetric formation of the passive domain. We note that the calculated extent of the quenched region obtained from using the velocity field is ∼ 5 *μ*m less than that obtained from using the intensity fluctuation fields - this difference could arise due to two reasons: first, from the asymmetric irregular domains that are present and have to be thresholded and second, from the differences between the variables being averaged (density fluctuations vs velocity fields). Nevertheless, the square root dependence on time approximately holds for both estimates. Since the azimuthally averaging process essentially ignores variations in motility inside the domain and its irregular shape, it is difficult to directly assign a physical mechanism behind the square root growth of the exposed domain.

In our experiments on the swarm, quenching due interaction between the bacteria and the light field effectively extracts actively generated energy of the swarm by degrading the ability of the bacteria to move and self-propel. If the mean velocity of the swarm arising due to the propagation of the swarm front is ignored, then one can treat the fluctuating swarm velocities, **v**, as time-varying fields with a zero mean when suitably averaged. The dyadic tensor **vv** then encodes information about the *energy content of the swarm*. Averaging over times scales long enough to encompass multiple vortex and streamer lifetimes, and then averaging azimuthally about the polar angle in the swarming plane, allows us to derive a radially dependent scalar field 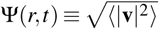 that encodes the time-averaged energy density. In principle, the evolution of this quantity will necessarily be coupled to the position dependent light field. While additional experiments are require to tease out the phenomenological form of the equation governing Ψ(*r,t*), it is interesting to compare the dynamics we observe with that in a much simpler, albeit similar physical problem - the one-dimensional freezing of a domain due to a heat sink at the origin that also yields a square root dependence in time for the boundary of the frozen domain (see also SI - §I).

### 3.3 Exposure time determines spatial extent of quenched domain

Next, we examine how varying the time of exposure while keeping the incident intensity field is fixed changes the size of the quenched domain. To adjust the viewing window, the 60 *μ*m aperture experiments are done with a 20x objective and the 120 *μ*m aperture with a 10x objective (we take into account the higher value for the intensity for the 60 *μ*m aperture case at 20x). To quantify the mean and variance of the data, we plot the average from four experiments with corresponding to the standard deviation shown as error bars. Aiming to obtain an upper bound on the extent of the swarming domain that is impacted, we choose to use intensity fluctuation fields rather than the PIV derived velocity fields.Tracing un-averaged boundary positions rj(t) from the image intensity fluctuations |Δ*I*\*| using Equation 2, we then estimate the effective size of the quenched region *r*_eff_ by calculating its radius of gyration following

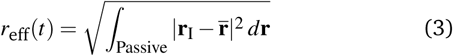

where 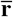 is the center of mass at time t. The maximum extent of the passive phase *R*_max_ is equal to *r*_eff_(*t* = *t*_off_); here the time at which exposure is terminated is *t*_off_.

From our experiments with two different aperture geometries, we find that *r*_max_ increases with *τ* and asymptotes to a constant that is slightly less than the aperture size (Fig. 5(c)); the asymptote is approximately 58 *μ*m for the 60 *μ*m aperture. For aperture size 120 *μ*m, we find the limiting asymptote to be ∼ 109 *μ*m. The data in Fig. 5(c) fits the functional form

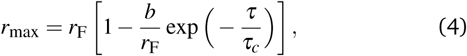

with *rF* = 58 *μ*m, *b =* 18 *μ*m, and *τ*_*c*_ = 33.5 s for the 60 *μ*m aperture and *r*_F_ = 109 *μ*m, *b* = 169 *μ*m, and *τ*_c_ = 37.3 s for the 120 *μ*m aperture.

### 3.4 Caging and jamming in quenched region

An unexpected feature seen in Fig 5(b) is that after exposure is terminated and the active bacteria invade and disrupt immotile bacteria, the size of the eroding region also roughly follow a square root dependence until the domain disperses completely. We examined if this feature was controlled by exposure time, *τ* (Figure 6a). The time complete dissolution, as expected, increases with exposure time. Surprisingly however, we find that the square-root scaling hold for both small as well as long exposure times. In our active, far-from equilibrium system, the passive domain is eroded by single bacteria-bacteria interactions (displacements originating from steric and self-propulsive mechanisms) as well as by collective highly non-equilibrium flow structures that form near the surface. Elucidating the origin of this scaling requires a consideration of the coupling between interface shape, interface speed and interface flow fields^33^.

**Fig. 6.**
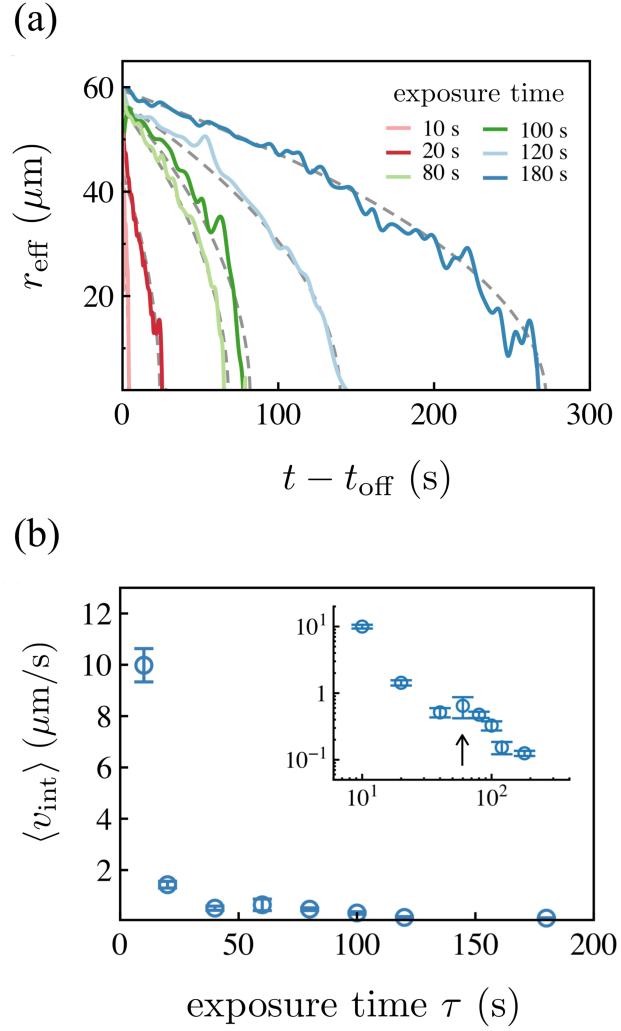
(a) The effective extent of the quenched, passive domain decreases over time *t* at rates that depend on the exposure duration *τ*. Longer exposure times prolong erosion by the active swarming bacteria, increasing the time it takes for the passive phase to disappear (at time *t*_0_). For each *τ*, the effective size *r*_eff_ follows 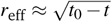 (grey dashed curves) with *t*_0_(*τ*) being the time for complete dissolution. (b) The average initial dissolution velocity (*v*_int_) decreases significantly with *τ*. Data is the average calculated from four experiments with standard deviation as error bars (Intensity ≈ 3 W/cm^2^).

Finally, we also measure the initial boundary dissolution speed by calculating ⟨*v*_int_⟩ = (*r*_eff_(*t*_off_) − *r*_eff_(*t*_off_ + Δ*t*))/ Δ*t*, where Δ*t* = 10 seconds. We find that ⟨*v*_int_⟩ varies significantly with exposure time - as shown in Fig 6(b) - decreasing from 10 *μ*m/s to 0.1 *μ*m/s as *τ* increases from 10 to 300 seconds. The decrease in speed is not monotonic, rather we observe a peak (inset) that is not ascribable to experimental variations. For long exposure times or high intensities, as the exposed bacteria are slowly paralyzed and slow down, they form jammed, highly aligned domains. Thus the time for complete erosion will be larger due to both the larger extent to be eroded as well as the aligned caged configurations of paralyzed bacteria.

## 4 Discussion and Perspectives

Previous work has highlighted the effect of light exposure - especially UV light on bio-molecular and biochemical mechanisms underlying propulsion in free swimming planktonic bacteria. Here, we presented work that complements these single cell experiments by analyzing the effects of light exposure on collective motility in swarming *Serratia marcescens*. In the absence of exposure, the swarming bacteria exhibit collective flows with significant vorticity (with a characteristic lifetime and frequency of formation) interspersed with streaming motions (with characteristic mean speeds)^33^. At low exposure levels, swarms are unaffected by light and maintain long-range collective motions. For sufficiently intense exposures, bacteria are rendered immobile and paralyzed, an effect that is either reversible or irreversible, depending on the exposure level. The permanently immobilized passive domain occurs for critical values of illumination power, requiring a minimum exposure time to appear. In the process, they form highly aligned, jammed domains whose size grows roughly as the square root of exposure time. Post exposure, active bacteria dislodge exposed bacteria from these caged configurations with initial dissolution rates strongly dependent on duration of exposure.

Interacting, high aspect ratio passive polar rods in the dense passive phase can organize into stable as well as metastable highly ordered states through thermal motions with additional steric interactions ^42^ or other weakly aligning interactions ^43^ - the degree of alignment governed by the relative strength of aligning interactions to the disordering random noise. Here, while activity leads to self-emergent collective flows, the slowing of individual bacteria, disruption of their self-propulsion and the further orienting affects of local shearing flows and tangentially acting steric interactions due to neighboring bacteria may allow the same mechanisms to operate resulting in the formation of strongly aligned and jammed phases.

Small regions of immobile bacteria can block space accessible to swarms, as well as trap motile bacteria and preventing them from escaping the light. Bacterial populations in nature include cells with a distribution of self-propulsion speeds. This raises the possibility that the subpopulations of exposed bacteria corresponding to the fastest moving cells or bacteria that are predisposed to UV resistance may escape from the exposed region. Reestablishing collective motility, and upon subsequent cell divisions, these cells may eventually result in emergence of resistant strains. Conversely, our experiments indicate that activity enhances alignment of neighboring cells. In collective swarms, these fast cells may surprisingly end up in jamming due to the preference to align. For bacteria with fixed propulsion and response at the organismal level, this implies that the time of exposure controls the fraction of cells immobilized as well as the degree of alignment. We thus hypothesize that the role of motility is subtle than is evident at first sight; and the exposure time is as important as light intensity in determining the degree of alignment in the exposed region. This in turn impacts the ability of the active bacteria to erode the boundary, enter the quenched region and dislodge the paralyzed cells.

Our experimental setup was suited to study the dynamics of the interphase region and the flows in its vicinity; thus we were able to study the time scales involved in the growth and initial erosion of the quenched region. Our experimental protocols cannot however examine a related interesting question - when can small sub-domains of bacteria can escape the exposed region before they are completely immobilized and trapped? It is intuitive to expect bacteria with low self-propulsion will be trapped; however, since the large scale complex flows are collective, adjacent adjacent that exhibit a velocity contrast can still move together due to the slow paralysis induced by light exposure and deactivation.

To complement our experimental observations, we propose and analyze a minimal (lumped) Brownian dynamics model to investigate the competition between propulsion speed and light-induced jamming underlying the ability of a test cell (mimicking a real cell-cluster) to escape from exposed regions. Working within the framework of an extended Langevin-like dynamics for the test cell (details in SI-§II), we assume that total light exposure increases the effective rotational diffusivity of the test cells; this may arise for instance from increased tumbling. At the same time, as the test cell moves in a mean field of its co-deactivating neighbors, its translational diffusion is hindered due to crowding and alignment. To include the effect of light on rotational diffusion, we turn to our experimental results and also previous studies on effects of light on *Bacillus subtilis* ^30^. Experiments show that that as bacterial cells become sluggish, the tendency to form flocks and large packs reduces and instead smaller clusters are observed. The overall reduction in cluster size and a less ordered motion within individual clusters gives rise to decreased correlation lengths with swarming eventually reverting to random motion in the presence of photodynamic effects. During exposure, the collective swarm velocity decays, a feature that can be recovered after exposure. Guided by these previous students and our experiments, we assume that light exposure increases the tumbling frequency and the rotational diffusion coefficient (the complementary case is also treated in the appendix). Note that bacteria also exhibit anisotropic translational diffusivities due to the different environments perpendicular and normal to the propulsion direction. Ignoring this anisotropy as well as changes to the self-propulsion speed (reduction in the self-propulsion speed alone is sufficient to immobilize the cell and so we do not pursue this), we find that the time evolution of trajectories is controlled by three dimensionless parameters: the Peclet number 𝒜_1_ charactering the self propulsion, and two deactivation parameters, 𝒜_2_ and 𝒜_3_, quantifying light induced changes in translational and rotational diffusion respectively. Salient results summarized in Figures 7 and 8 show that the Peclet number controls how far particles can travel before losing mobility in an unbounded as well as bounded light fields. We are currently studying the quenching process using detailed agent based simulations that account for the rod geometry of the bacteria, hydrodynamic interactions by modeling the bacteria as force dipoles and excluded volume interactions via the Maier-Saupe potential. These studies will also shed light on the mixing of paralyzed bacteria in the active region.

**Fig. 7.**
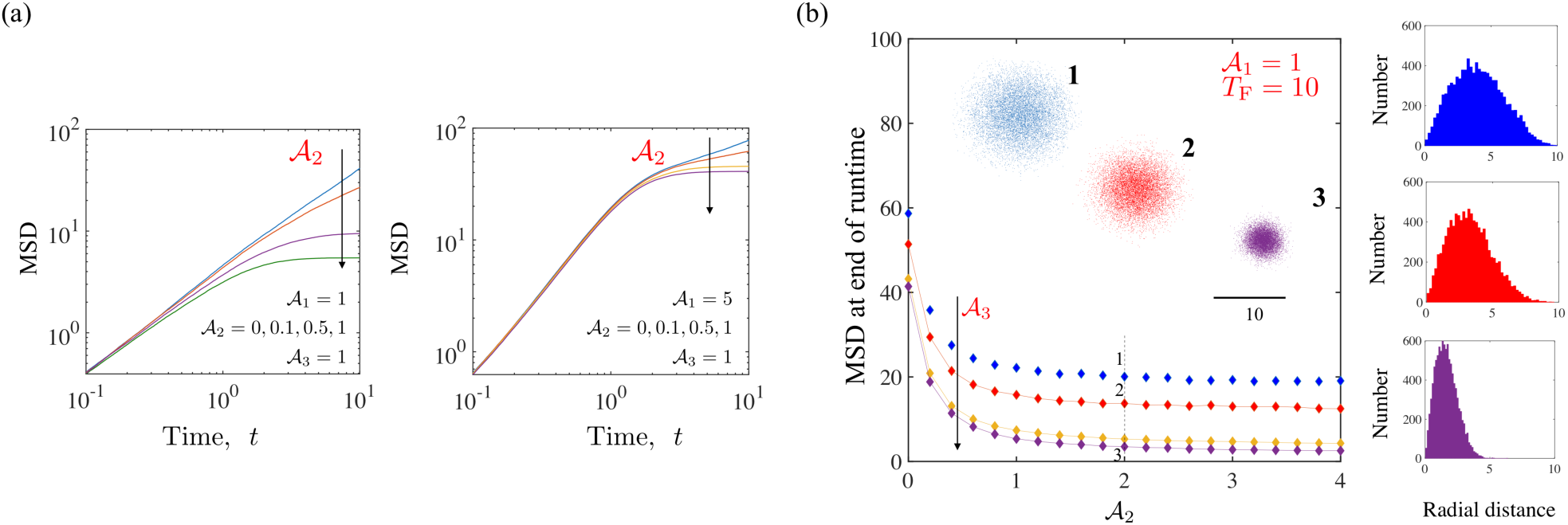
Dynamics of self-propelled and diffusing particles (*N* = 10^4^) interacting with a constant, unbounded light field (Φ (**r**) = 1, *R*_L_ = ∞). Light exposure modifies the translational and rotational diffusivities, but not the self-propulsion speed. Cells are released at the origin *r* = 0 and trajectories integrated for a dimensionless time interval *T*_F_ = 10 with Δt ∈ (4 × 10^—4^,10^—3^). (a) Examination of the ensemble averaged MSD(*t*) shows that trajectories becoming denser and compact yielding a plateau in the MSD corresponding to localization and paralysis of the particles. Changes in rotational diffusivities are required for this to happen since the self-propulsion speed is assumed to be constant; this effect is exacerbated as 𝒜_2_ becomes larger. Note that as 𝒜_1_ increases, the longer the particles typically travel before exposure effects dominate. (b) MSD (*t* = *T*_F_) as a function of 𝒜_2_ for various values of 𝒜_3_ (from top to bottom: 0.1, 0.5, 1 and 2) with 𝒜_1_ = 1. Consistent with (a), we observe saturation for 𝒜_2_ > 3. (Inset) Shown are the (*x,y*) locations of the particles for parameters corresponding to points 1, 2 and 3 marked on the plot at *T*_F_ = 10. Examination of the corresponding number distribution plots (right tiles) shows the peak shifting to lower values of radial distance r, and significant changes to the tail end of the distribution function. Since the light field is unbounded, all particles are eventually affected. For particles with low Peclet number (low activity), the exposure time determines how far they can travel before becoming deactivated.

**Fig. 8.**
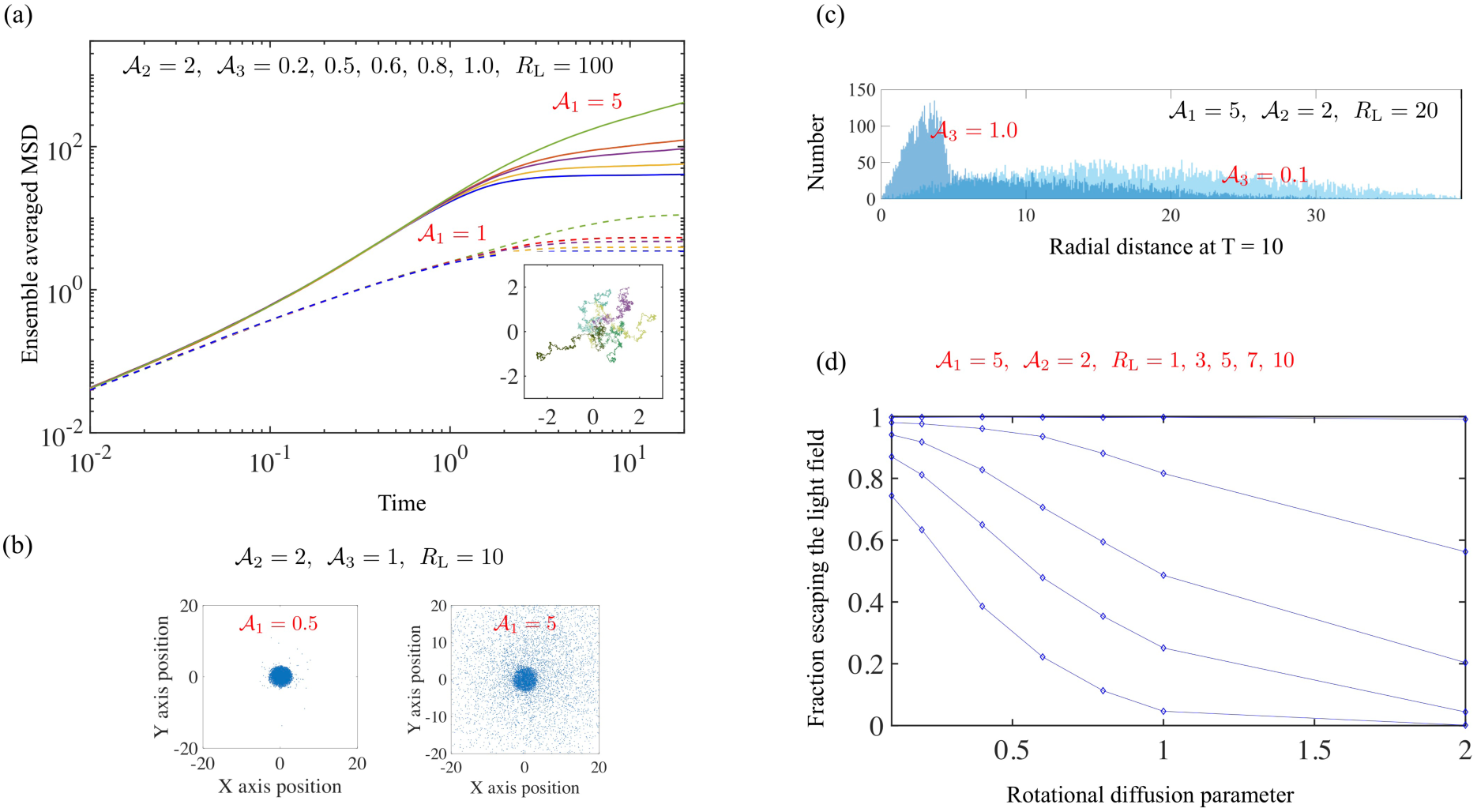
Effects of an imposed (dimensionless) length scale *R*_L_ using a finite extent light field F(**r**) = 1 Vr e (0,R_L_) and zero elsewhere. We integrate trajectories of 10^4^ particles using Δt = 4 × 10^—4^ up to final times *T*_F_. (a) First effects of finite light extent. The dimensionless MSD is shown as a function of time for *R*_L_ = 100. Solid curves are results for 𝒜_*1*_ = 5 while dashed curves are for 𝒜_1_ = 1. We see that as 𝒜_3_ increases from 0.2 to 1.0, MSD saturates rapidly. (Inset) Sample trajectories for 𝒜_1_ = 1 for 0 < *t* < 20 demonstrating localization for *t* > 6. (b) Decreasing from 100 to 10 brings out the role of Peclet number in enabling escape. We show cell positions at *t* = *T*_F_ = 10 (note that cells outside of this domain are not shown). (c) The fraction of 10^4^ trajectories that start at *r* = 0 and are located at *r* > *R*_L_ at *T* = 10. Note that some of these trajectories may end up reentering the domain in the simulation but not in the experiment due to the dense, jammed domains that prevent reentry. The curves are shown as a function of the rotation diffusion parameter 𝒜_3_.

A natural extension of our experiments would be to systematically isolate the effects of light on aspects of cell-cell interaction and communication that determine swarm survival and virulence including individual cell motility, collective motion and active/passive particle interactions^44^. Experiments using filters that allow for wavelength dependent immobilization at the single cell level will allow us to further understand how to collective motility and light exposure are related.

## Acknowledgements

We thank Ed Steager and Elizabeth Hunter for providing the cells and for experimental assistance. AEP was supported by an NSF Graduate Research Fellowship. PEA acknowledges funds from NSF-DMR-1104705 and NSF-CBET-1437482. AG acknowledges startup funds from the University of California, Merced and funding from UC 2018 Senate Award program.

## Author Contributions

AEP, PA and AG conceived the study. AEP and AG designed and performed the experiments, analyzed the data and wrote the manuscript. AG designed the simulation model and analyzed results. JY performed simulations of the model. All authors gave final approval for submission.

## Data availability

The authors declare no competing financial interests. Correspondence and requests for materials should be addressed to AG or to AEP. All relevant data are available from the authors upon request.

## References

1. L. Alberti and R. M. Harshey, Journal of bacteriology, 1990, 172, 4322.

2. R. M. Harshey, Annual Reviews in Microbiology, 2003, 57, 249.

3. N. Verstraeten, Trends in microbiology, 2008, 16, 496.

4. M. F. Copeland and D. B. Weibel, Soft matter, 2009, 5, 1174.

5. N. C. Darnton et al. Biophysical journal, 2010, 98, 2082.

6. L. Turner, R. Zhang, N. C. Darnton and H. C.. Berg, Journal of bacteriology, 2010, 192, 3259.

7. D. B. Kearns, Nat. Rev. Microbiol., 2010, 8, 634.

8. R. M. Harshey and J. D. Patridge, Journal of molecular biology, 2015, 427, 3683.

9. E. B. Steager, C. B. Kim and M. G. Kim, Physics of Fluids, 2008, 20, 073601.

10. M. T. Butler, Q. Wang and R. M. Harshey, Proceedings of the National Academy of Sciences., 2010, 8, 3776.

11. D. Roth et al, Environmental microbiology, 2013, 15, 2532.

12. S. Lai, J. Trembley and E. Deziel, Environmental microbiology, 2009, 11, 126.

13. C. Chen, S. Liu, X. Shi, H. Chate and Y. Wu, Nature, 2017, 542, 210.

14. H. C. Berg and D. A. Brown, Nature, 1972, 239, 500.

15. H. C. Berg, E. coli in Motion. 2008, Springer.

16. G. H. Wadhams and J. P. Armitage, Nat. Rev. Mol. Cell Biol., 2004, 5, 1024.

17. H. C. Berg and R. A. Anderson, Nature, 1973, 245, 380.

18. A. E. Patteson, A. Gopinath, M. Goulian and P. E. Arratia, Scientific Reports, 2015, 5, 15761.

19. B. L. Taylor and D. E. Koshland Jr., Journal of bacteriology, 1975, 123, 557.

20. S. Wright et. al., Journal of bacteriology, 2006, 188, 3961.

21. E. Steager et. al., Applied Physics Letters, 2007, 90, 26.

22. M. P. Conley and H. C. Berg, Journal of bacteriology, 1984, 158, 832.

23. V. Kodoth and M. Jones, http://medicalmate.gr/img/cms/UVC, 2015, 1.

24. L. Alonso-Saez, J. M. Gasol, T. Lefort, J. Hofer and R. Sommaruga, Appl Environ Microbiol., 2006, 72(9), 5806.

25. D. G. Sharp, J. Bacteriol., 1940, 39(5), 535.

26. M. B. Said, S. Khefacha, L. Maalej, I. Daly and A. Hassen, African J. of Microbiol. Res., 2011, 5(25), 4353.

27. B. Li and B. E. Logan, Colloids Surf B Biointerfaces, 2005, 41(2-3), 153.

28. C. M. Abana et. al., Microbiologyopen., 2017, 6(4), e00466.

29. T. Dai, M. S. Vrahas, C. K. Murray and M. R. Hamblin, Expert review of anti-infective therapy, 2012, 10, 185.

30. S. Lu, W. Bi, X. Wu, B. Xing and E. K. L. Yeow, Phys. Rev. Lett., 2013, 111, 208101.

31. T. Dai et. al., Antimicrobial agents and chemotherapy, 2013, 57, 1238.

32. S. Pei, A. C. Inamadar, K. A. Adya and M. M. Tsoukas, Indian Dermatol Online J., 2015, 6, 145. 10

33. A. E. Patteson, A. Gopinath and P. E. Arratia, Nature Communications, 2018, 9, Article number: 5373.

34. A. E. Patteson, A. Gopinath and P. E. Arratia, Current Opinion in Colloid & Interface Science, 2016, 21, 86.

35. S. Bensity E. Ben-Jacob, G. Ariel and A. Be’er, Phy. Rev. Lett., 2015, 114, 018105.

36. W. Thielicke and E. Stamhuis, Journal of Open Research Software, 2014, 2, 1.

37. http://zeiss-campus.magnet.fsu.edu/print/lightsources/mercuryarc-print.html

38. Y. Wu and H. C. Berg, Proc. Nat. Acad. Sci., 2011, 109, 4128.

39. A. Gopinath, R. C. Armstrong and R. A. Brown, J. Cryst. Growth, 2006, 291(1), 272.

40. R. Cerbino and V. Trappe, Physical Review E, 2008, 80, 031403.

41. L. G. Wilson et. al., Phys. Rev. Lett., 2011, 106, 018101.

42. A. Gopinath, R. C. Armstrong and R. A. Brown, J. Chem. Phys., 2004, 18, 028102.

43. A. Gopinath, L. Mahadevan and R. C. Armstrong, Phys. Fluids, 2006, 121 (12), 6093.

44. A. E. Patteson, A. Gopinath, P. K. Purohit and P. E. Arratia, Soft Matter, 2016, 12(8), 2365.

45. H. S. Carslaw and J. C. Jaeger, Conduction of heat in solids, 1959, Oxford Science, 2nd edition.

46. G. Barenblatt, Scaling, Self-similarity, and Intermediate Asymptotics: Dimensional Analysis and Intermediate Asymptotics, 1996, Cambridge University Press.

47. G. Volpe, S. Gigan and G. Volpe, Am. J. Physics, 2014, 82, 659.

